# A human monoclonal antibody from hybrid immunity exhibits exceptional neutralization breadth against antigenically divergent contemporary Omicron JN.1 lineage variants

**DOI:** 10.64898/2026.07.20.739483

**Authors:** Diwakar Rathour, Shweta Shrivas, Naresh Kumar, Chaman Prasad, Janmejay Singh, Gagandeep Singh, A. Surendranath, Shweta Rathi, Sneha Verma, Supriya Chauhan, Balwant Singh, Jyoti Sutar, Souvick Chattopadhyay, Gaurav Batra, Sudipta Sonar, Isabella Eckerle, Lorenzo Subbissi, Anant Mohan, Shailendra Asthana, Suprit Deshpande, Jayanta Bhattacharya

**Affiliations:** Center for Virus Research, Therapeutics & Vaccines, BRIC-Translational Health Science & Technology Institute, NCR Bioscience Cluster, Faridabad, Haryana-121001, India; Monoclonal Antibody Biofoundry Unit, BRIC-Translational Health Science & Technology Institute, NCR Bioscience Cluster, Faridabad, Haryana-121001, India; Bioassay Laboratory, BRIC-Translational Health Science & Technology Institute, NCR Bioscience Cluster, Faridabad, Haryana-121001, India; Computational Biophysics and Computer Aided Drug Discovery group, Center for Computational & Mathematical Biology, BRIC-Translational Health Science & Technology Institute, NCR Bioscience Cluster, Faridabad, Haryana-121001, India; Department of Pulmonary, Critical Care and Sleep Medicine, All India Institute of Medical Sciences, New Delhi- 110029, India; Center for Biodesign & Diagnostics, BRIC-Translational Health Science & Technology Institute, NCR Bioscience Cluster, Faridabad, Haryana-121001, India; Geneva Centre for Emerging Viral Diseases, University Hospitals of Geneva and Medical Faculty, University of Geneva, Switzerland; Department of Medicine, Medical Faculty, University of Geneva, Geneva, Switzerland; Department of Microbiology and Molecular Medicine, University of Geneva, Geneva, Switzerland; World Health Organization, Geneva, Switzerland; Antibody Translational Research Program, BRIC-Translational Health Science & Technology Institute, NCR Bioscience Cluster, Faridabad, Haryana-121001, India

## Abstract

Hybrid immunity offers stronger and more durable antibody-mediated protection against symptomatic SARS-CoV-2 infection. In the present study, we report the development of a durable antibody response in an individual with hybrid immunity who also received three doses of the prototype COVID-19 vaccine. Polyclonal plasma antibody obtained from this donor also showed extraordinary neutralization breadth against contemporary Omicron variants. One of the functional monoclonal antibodies (ATHS-C30) isolated from this individual, representing the IGHV4-30 lineage-specific B cell with strong binding affinity to JN.1 spike protein, showed extraordinary neutralization breadth, including variants of Omicron lineages that emerged beyond JN.1, such as KP.2, KP.3.1.1, KP.3.2, KP.3.3, LB.1 and XEC. ATHS-C30 was found to bind RBD with high affinity and showed distinct epitope specificity to the other neutralizing mAbs isolated from the same donor through the epitope binning assay. Molecular modelling of the CDHR3 sequence using existing structures indicated that ATHSC-30 belongs to the class 4 antibody, a feature that contributes to breadth, while epitope conservation analysis indicated that the majority of RBD-interacting residues of ATHSC-30 are evolutionarily conserved. Taken together, our study indicate that ATHS-C30 forms the basis of development of a robust and broadly neutralizing antibody response in this individual with hybrid immunity, which overcomes the antigenic variation by targeting highly conserved and cryptic epitopes in destabilizing the spike structure.

**Importance:** SARS-CoV-2 continues to pose a significant public health threat, particularly to immunocompromised individuals and older adults with underlying comorbidities. Hybrid immunity to SARS-CoV-2 in vaccinated individuals leads to the development of B cells that are qualitatively superior to those that are expected to develop in only vaccinated individuals. In the present study, we found that among individuals with hybrid immunity who developed robust, durable antigen-specific antibody responses, one developed antibody response capable of broadly cross-neutralising contemporary Omicron variants. This was correlated with development of antigen-specific B cell lineage, such as IGHV4-30, that produced antibodies with potent and extraordinary neutralisation breadth against contemporary Omicron lineages, such as KP.2, KP.3.1.1, KP.3.2, KP.3.3, LB.1 and XEC. This is believed to be due to heterologous antigen exposures driving the development of an antigen-specific B cell repertoire that, in turn, facilitates immune imprinting capable of overcoming the ineffectiveness of antibodies to effectively neutralize newly emerging Omicron variants.

## Introduction

Although the global burden of COVID-19 has declined, the continued evolution of SARS-CoV-2 remains a significant challenge, particularly for immunocompromised individuals and the elderly with underlying comorbidities, who are at increased risk of severe disease and may derive suboptimal protection from vaccination. The pandemic landscape of SARS-CoV-2 has been drastically altered by its continued evolution from massive infection surges during the initial phase of the pandemic to becoming endemic over time, accompanied by the frequent emergence of Omicron lineage subvariants, which are known for their adaptability and immune evasion, leading to milder symptoms, especially in vaccinated populations with hybrid immunity (1–5). The emerging Omicron lineage variants with complex mutations accumulated over time in the receptor binding domain (RBD) of the spike protein that disrupts access to the neutralizing antibodies pose concerns for vaccine and monoclonal antibody (mAb) mediated effectiveness (6–8). While the durability of the effectiveness of the ancestral Wuhan strain-based vaccine immunogen has been challenged by the emergence of heavily mutated variants, hybrid immunity has been shown to provide the strongest protective immunity against evolving variants particularly in protecting against severe disease outcomes (9–11). Moreover, booster doses of existing vaccine immunogens have shown that they remain beneficial for protecting against moderate and severe disease, regardless of the initial course (3, 12). It is believed that the broadly protective immunity observed in individuals with hybrid immunity is due to the higher quality of the antibody response they mount compared to that elicited by vaccination alone (13, 14). This has been supported by the isolation of several broadly neutralizing monoclonal antibodies (mAbs) from such individuals (15–26). Several of these mAbs were also used in clinical settings to prevent disease exacerbation in symptomatic patients due to their qualitative protective and therapeutic properties (17, 18, 20, 21).

In the present study, we examined magnitude, durability and neutralization specificity of antibodies developed in longitudinally followed-up individuals with a history of infection and who received the initial two doses of the prototype vaccine (BBV152 or ChAdOx1nCoV-19) with a third booster vaccine dose. Amongst them, we identified one donor (A0015), who received three doses of ChAdOx1nCoV-19 and developed antibodies with strong specificity to JN.1 and demonstrated neutralization breadth across contemporary Omicron lineage variants. mAbs isolated from this individual revealed SARS-CoV-2 specific B cell lineages that showed variable neutralization against Omicron variants including one (ATHS-C30) that showed potent and extraordinary neutralization breadth of contemporary Omicron variants including JN.1, KP.2, KP.3.1.1, KP.3.2, KP.3.3, LB.1 and XEC. While these observations form the basis for the development of robust humoral immunity in this particular individual with broad antigen specificity, mAbs such as this one (ATHS-C30) can also be harnessed to develop pan-SARS-CoV-2 vaccine candidates, in addition to their potential for direct therapeutic applications.

## Results

### Identification of an individual with hybrid immunity who mounted durable humoral immunity of high magnitude against contemporary SARS-CoV-2 circulating Omicron variants

We examined a total of 26 individuals with hybrid immunity to SARS-CoV-2 and who received three doses of vaccine (Table S1) to first assess (a) the magnitude of the antibody response developed against the ancestral SARS-CoV-2 RBD (which was part of the composition of all the vaccines manufactured and taken by all the donors) and (b) their neutralization breadth across Omicron variants known to evade most neutralizing antibodies elicited by vaccines and several monoclonal antibodies. Plasma antibodies obtained from these individuals between 6-12 weeks after receiving the third booster dose of the vaccine (Table S1) were first assessed by quantitative RBD ELISA for magnitude, followed by their durability over the next three months. As shown in **Figure 1A**, we found that all the individuals examined developed a robust ancestral RBD-specific antibody response with comparable antibody titers of 550.72 and 595.97 BAU/mL, with plasma samples obtained at baseline and after 90 days, respectively, indicating durability. Next, we examined the neutralization profiles of the plasma antibodies against contemporary Omicron variants by pseudovirus neutralization assay. We observed that while plasma antibodies developed in the majority of the donors (>90%) collected at the baseline (post receiving third vaccine dose) showed strong neutralization of BA.1 and BA.2.75, including ancestral and Delta variants, only few of them showed neutralization (those that demonstrated >60% neutralization at 1:100 plasma dilutions) of pseudoviruses expressing BQ.1.1 (8/26), HV.1 (2/26), XBB1.5 (8/26) and JN.1 (1/26) spikes (**Figure 1B**). Interestingly, while no neutralization of XBB.1.5 and JN.1 by plasma antibodies obtained at the baseline in 5/26 donors (A0023, A0025, A0026, A0030, A0032) was observed, we found that plasma antibodies obtained at the second visit from these donors were capable of strongly neutralizing XBB1.5 (>80% neutralization observed with 1:100 dilution of plasma) with visit 2 plasma obtained from A0032 donor also showed >80% neutralization of JN.1 compared to no neutralization observed with the plasma sample obtained at the baseline. This observation indicated that these individuals were likely exposed to XBB.1.5. Amongst all, one of the donors (A0015) was found to develop antibodies that showed broad and potent neutralization of the Omicron variants tested, including JN.1. We next examined the extent of binding of the plasma antibodies obtained from A0015 donor at the baseline to both ancestral and JN.1 RBDs at different plasma dilutions. It indeed bound both the ancestral and JN.1 RBD proteins in a dose-dependent manner (**Figure 2A**). Subsequently, a dose-response neutralization assessment of A0015 plasma antibodies against a panel of Omicron variants was next carried out. As shown in **Figure 2B**, A0015 plasma antibodies neutralized pseudoviruses expressing BA.1, BA.2, BA.4, B.2.75, BQ1.1, HV.1, XBB.1.5 and JN.1 with >90% neutralization found against BA.1, BA.2, BA.4, and B.2.75 by plasma antibodies diluted at 1:540 and >80% neutralization observed against BQ.1.1, HV.1 and JN.1 with same at 1:180 dilution. Furthermore, A0015 plasma antibodies also showed cross-neutralization of KP.2, KP.3.1.1, KP.3.2, KP.3.3, LB.1 and XEC Omicron variants besides XBB.1.6 and JN.1 by live authentic virus neutralization assay (**Figure 2C**). Taken together, we identified A0015 as an individual with hybrid immunity who developed a robust antibody response with a higher order of magnitude and capable of broadly neutralizing contemporary Omicron variants representing distinct lineages.

**Figure 1.**
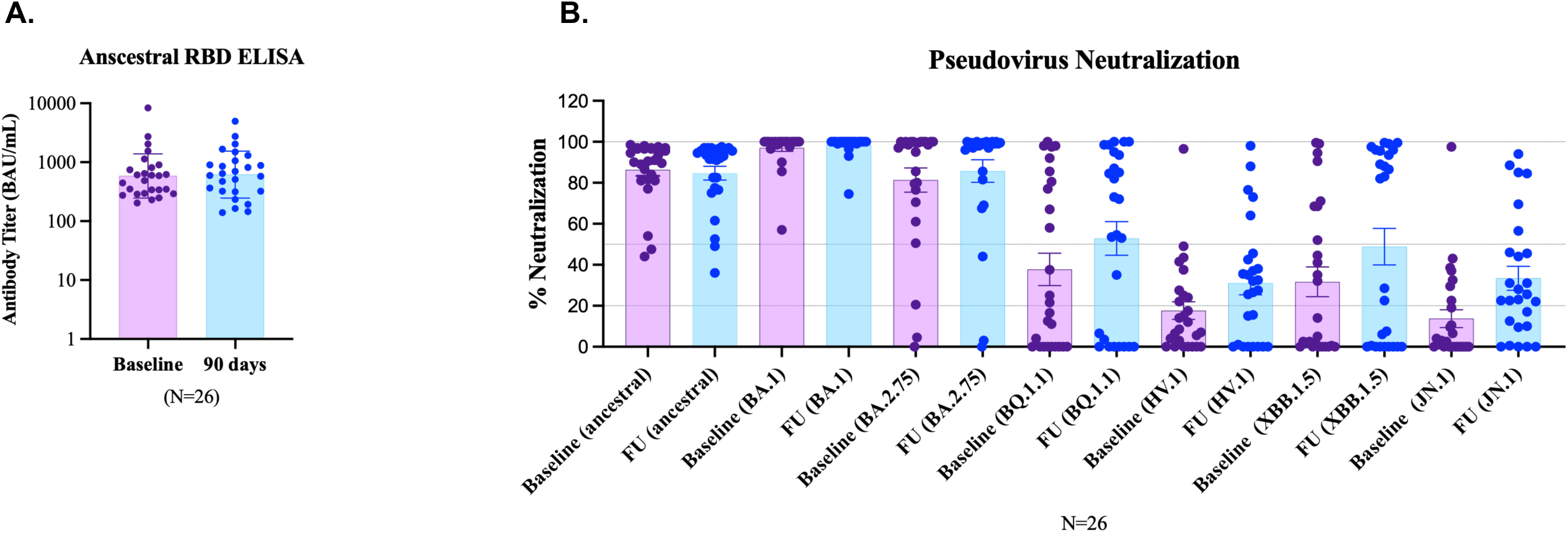
Dynamics of SARS-CoV-2-specific antibody response in individuals (N=26) with hybrid immunity. **A.** Magnitude and durability of total ancestral RBD-specific antibodies post-receiving third booster vaccine dose as determined by quantitative dose-response RBD ELISA; **B.** Neutralization profile of plasma antibodies against Omicron variants. Pseudovirus neutralization assay was carried out using 1:100 diluted heat-inactivated plasma samples as a source of antibodies developed in these individuals.

**Figure 2.**
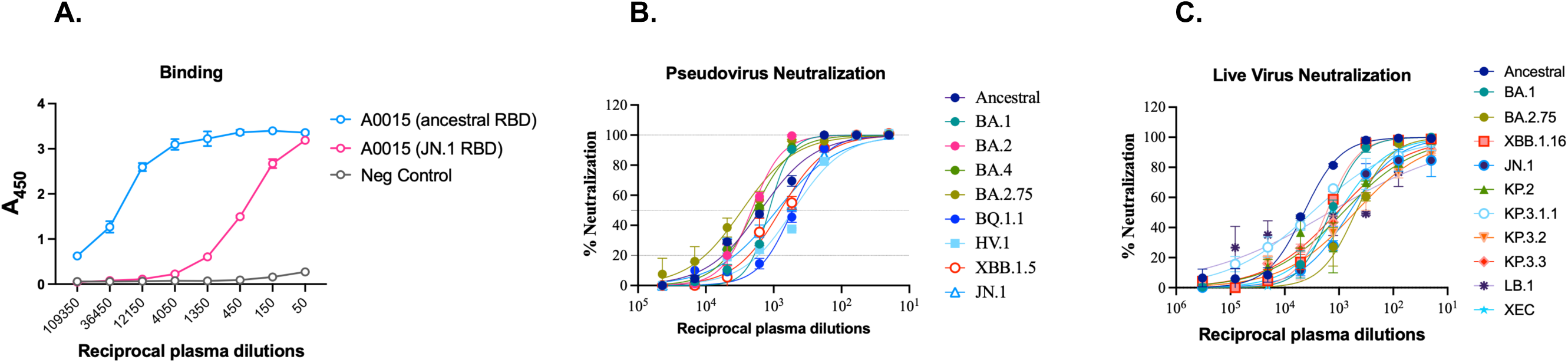
Binding and neutralizing antibodies developed in A0015 donor. Dose-response binding of plasma antibodies obtained from A0015 baseline to ancestral and JN.1 RBD soluble proteins as determined by ELISA. Heat-inactivated plasma from a healthy individual obtained prior to 2019 was used as negative control **(A)**. Neutralization profile of A0015 plasma antibodies to contemporary Omicron variants was determined by pseudovirus (**B)** and live neutralization assay (**C)** respectively. A0015 plasma was found to neutralize all the Omicron variants in a dose-dependent manner.

### Isolation of a monoclonal antibody from A0015 donor capable of neutralizing JN.1

To delineate the basis of substantial neutralization breadth by antibodies developed in A0015 donor, we next isolated RBD-specific monoclonal antibodies (mAbs) from this particular donor by antigen-specific single B cell sorting as described before (27). Biotinylated ancestral Wuhan and JN.1 RBD were used as antigen baits to singly sort IgG^+^/IgM^-^/IgD^-^/CD19^+^/CD20^+^/CD3^-^/CD14^-^/CD16^-^/CD8^-^ B cells in 96-well plate (**Figure S1A**). mRNA from the lysed single B cells from each well of the 96-well plate were used as template to amplify variable heavy and light chain IgG sequences. Paired variable heavy and light chain IgG sequences were subsequently cloned into expression vectors and recombinant mAbs expressed in HEK293T cells. Cell supernatants expressing recombinant mAb clones were further screened to identify functional mAb clones reactive to ancestral Wuhan and JN.1 RBD proteins. 11/26 (42.3%) mAb clones were found to be strongly reactive to ancestral Wuhan RBD, as shown in ELISA binding, out of which one ( ATHS-C30) also showed binding to JN.1 RBD (**Figure S1B**). In the preliminary screening using cell supernatants, out of the nine antigen-reactive mAb clones, five (ATHS-C24, ATHS-C30, ATHS-C34, ATHS-C48, and ATHS-C52) were found to show strong neutralization of pseudovirus expressing ancestral SARS-CoV-2 spike protein of which ATHS-C30 also showed neutralization of pseudovirus expressing JN.1 that correlated with its strong binding to JN.1 RBD as shown above and as expected (**Table 1**).

**Table 1.** Screening of cell supernatants representing ancestral RBD-positive A0015 B cell clones for their neutralization potential.

| Serial no | Clone ID | Pseudovirus neutralization* |  |
| --- | --- | --- | --- |
|  |  | Ancestral | JN.1 |
| 1 | ATHS-C09 | + | - |
| 2 | ATHS-C11 | ++ | - |
| 3 | ATHS-C15 | + | - |
| 4 | ATHS-C21 | + | - |
| 5 | ATHS-C22 | + | - |
| 6 | ATHS-C24 | ++++ | - |
| 7 | ATHS-C30 | ++++ | +++ |
| 8 | ATHS-C34 | ++++ | - |
| 9 | ATHS-C36 | ++ | - |
| 10 | ATHS-C40 | ++ | - |
| 11 | ATHS-C43 | ++ | - |
| 12 | ATHS-C46 | +++ | - |
| 13 | ATHS-C47 | + | - |
| 14 | ATHS-C48 | ++++ | + |
| 15 | ATHS-C52 | ++++ | - |
| 16 | ATHS-C56 | ++ | - |
| 17 | ATHS-C60 | ++ | - |
| 18 | ATHS-C64 | ++ | - |
| 19 | ATHS-C71 | ++ | - |
| 20 | ATHS-C74 | ++ | - |
| 21 | ATHS-C86 | + | - |
| 22 | ATHS-C98 | ++ | - |
| 23 | A0015 plasma | ++++ | +++ |

\* Note that neat cell supernatants were used for the neutralization

The variable heavy and light chain IgG sequences of these five mAb clones revealed their unique B cell germline origin, with heavy variable IgG chain CDHR3 length varied from 16 to 21 and variable light chain IgG CDR3 from 9 to 10 amino acids respectively (**Table 2**). Between 90.28% and 96.14% (variable heavy chain) and 91.04% and 98.58% (variable light chain) were identical to the respective germlines (**Table 2**), indicating minimal somatic mutations in the newly isolated mAbs. Taken together, we isolated five mAbs from the A0015 donor by B cell cloning, of which one (ATHS-C30) was found to be capable of binding and neutralizing JN.1 RBD and pseudovirus expressing JN.1 spike.

**Table 2.**
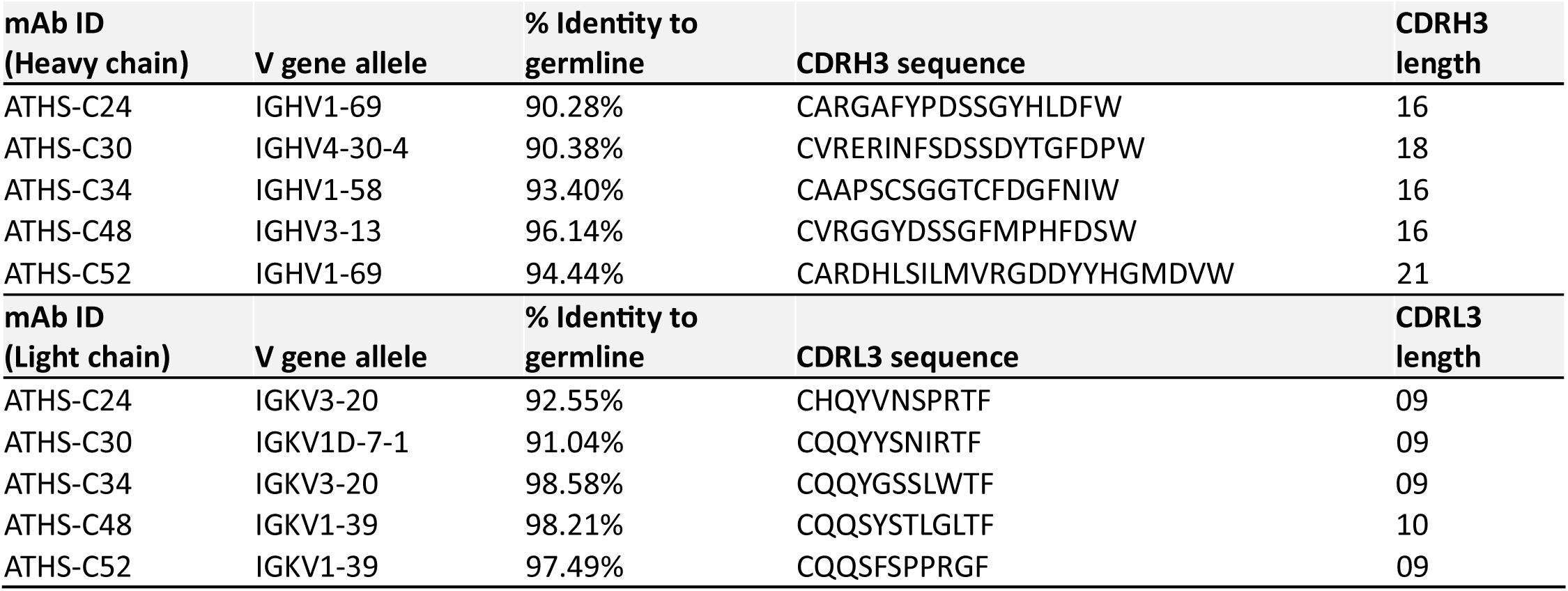
Genetic feature of variable heavy and light chain IgGs of novel mAbs.

### ATHS-C30 demonstrated strong RBD binding, stability and exceptional neutralization breadth across contemporary Omicron variants

The five novel mAbs isolated from the A0015 donor, purified as IgG, were next examined for their binding to both ancestral and JN.1 spike proteins by ELISA. While all the purified mAbs showed comparable binding to ancestral RBD protein (**Figure 3A**), only ATHS-C30 showed binding to the JN.1 RBD in ELISA in a dose-dependent manner (**Figure 3B**), consistent with observations made above with the cell supernatant from transfected 293T cells. This observation further confirmed the exclusivity of ATHS-C30 as the unique mAb amongst the others to bind to the JN.1 RBD. A stronger binding of ATHS-C30 to the ancestral RBD (EC_50_ of 0.009 µg/mL) than to the JN.1 RBD (EC_50_ of 0.32 µg/mL), however, was observed. A similar observation was made when we next examined the extent of binding to both ancestral and JN.1 spikes expressed on the cell surface of 293T cells. As shown in **Figure 3C**, compared to other mAbs, ATHS-C30 showed dose-dependent binding to JN.1 spikes expressed on the cell surface, further indicating its ability to interact with functionally active ancestral Wuhan and JN.1 spike protein expressed in their native forms. ATHS-C20 also showed binding kinetics similar to those of ATHS-C30 to both ancestral and JN.1 RBDs, as observed in the BLI analysis, consistent with ELISA binding. As shown in **Figure 3D**, ATHSC30 demonstrated stronger binding to the ancestral RBD (K_D_ of <1 x 10^-12^ M) compared to ATHS-C24, ATHS-C34, ATHS-C48, and ATHS-C52 (K_D_ ranged from 1 x 10^-12^ to 9.9 x10^-11^ M; **Figure S2**). Furthermore, ATHS-C30 was also found to bind soluble JN.1 trimeric spike protein with moderate affinity, with a K_D_ of 3.26 × 10^-12^ M with X and Y as its K_on_ and K_off_ rates, respectively (**Figure 3E**). Overall, based on the above observations, we concluded that ATHS-C30 binds to functional JN.1 spikes with high specificity.

**Figure 3.**
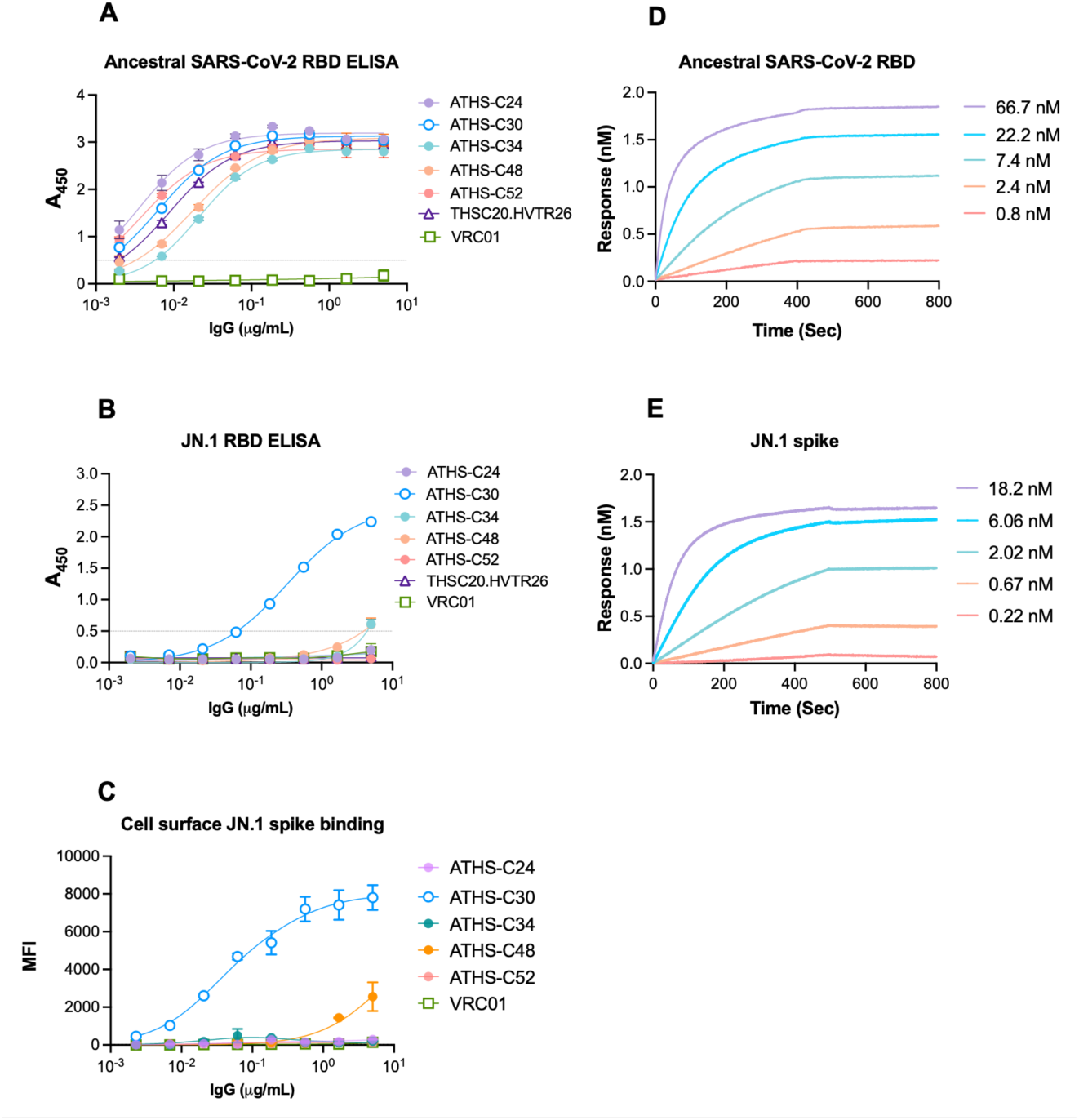
Binding of novel mAbs isolated from A0015 donor to ancestral and JN.1 RBD. ELISA binding of ATHS-C24, ATHS-C30, ATHS-C34, ATHS-C48 and ATHS-C52 to ancestral **(A)** and JN.1 RBD **(B)**. Note that while ATHS-C30 bound to JN.1 RBD in a dose-dependent manner, none of the other autologous mAbs isolated from A0015 bound to JN.1 RBD. **C.** Binding of the ATHS-C30 to JN.1 spike trimers in native form expressed on the surface of 293T cells was assessed by flow cytometry. Note that other ATHS-C30, none of the other mAbs bound to JN.1 spike trimers on the cell surface. Binding kinetics of ATHS-C30 to ancestral RBD **(D)** and JN.1 spike **(E)** protein as assessed by BLI.

Next, we examined the extent of neutralization breadth of purified ATHS-C30 IgG against a panel of Omicron variants (including those that emerged after JN.1 outbreak) both by pseudovirus and authentic live virus neutralization assays. As shown in **Figure 4A**, ATHS-C30 showed potent neutralization of pseudoviruses expressing ancestral SARS-CoV-2, Delta, BA.1, BA.2, BA.4, BA.2.75, BQ.1, HV.1, XBB.1.5 and JN.1 spikes with IC_50_ of 0.07, 0.07, 0.04, 0.05, 0.06, 0.09, 0.04, 0.05, 0.02 and 0.05 μg/mL, respectively. In addition, ATHS-C30 was also found to show potent neutralization of BA.5, XBB.1.16, JN.1, KP.2, KP.3.1.1, KP.3.2 , KP.3.3, LB.1 and XEC replication competent isolates with the high potency of 0.054, 0.044, 0.151, 0.067, 0.028, 0.035, 0.04, 0.205 and 0.018 μg/mL respectively in live virus neutralization assay (**Figure 4B**). Taken together, our data indicate ATHS-C30 as a unique broadly neutralizing novel mAb isolated from the A0015 donor that showed potent neutralisation of all the contemporary Omicron variants tested in this study. It also highlights its contribution to the exceptional neutralization breadth with higher magnitude mounted in this particular donor with hybrid immunity, capable of overcoming the evolving mutational constraints accumulated in the spike protein that are correlates of evasion of evolving variants to existing neutralizing antibodies.

**Figure 4.**
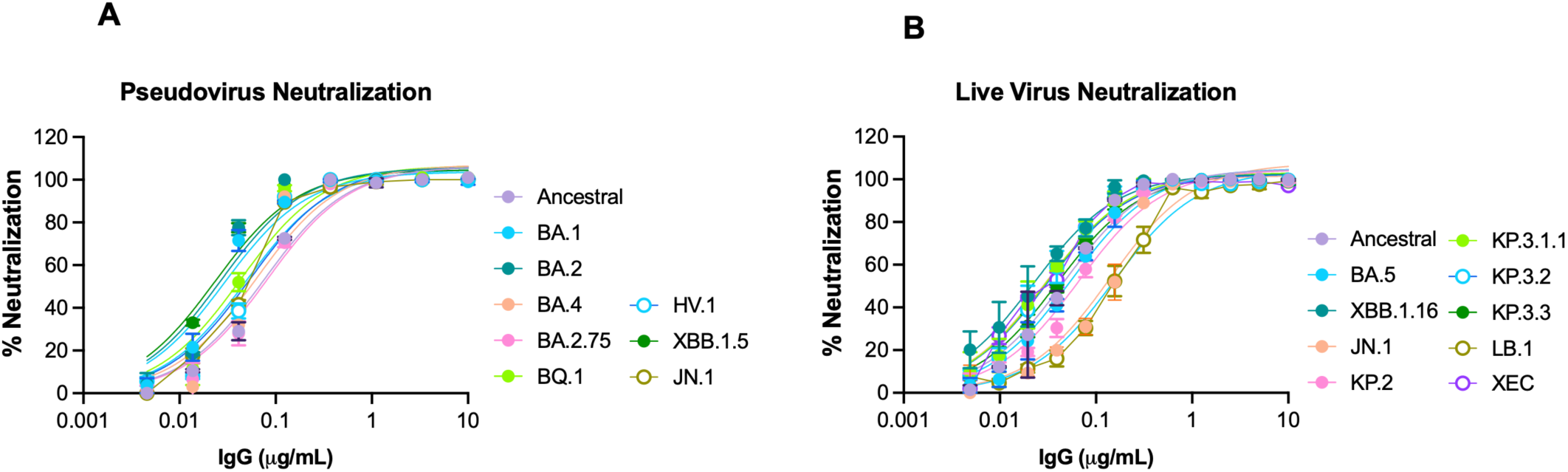
Neutralization breadth of ATHS-C30. Pseudovirus neutralization by ATHS-C30 of BA.1, BA.2, BA.4, BA.2.75, BQ.1, HV.1, XBB.1.5 and JN.1 was carried out in 293T cells overexpressing ACE-2 (**A)** and live authentic virus neutralization of BA.1, BA.5, XBB.1.16, JN.1, KP.2, KP.3.1.1, KP.3.2, KP.3.3, LB.1 and XEC by ATHS-C30 was assessed in Vero-E6 cells. IC_50_ values were calculated using GraphPad Prism software.

### ATHS-C30 targets non-competing epitopes on RBD compared to neutralizing mAbs isolated from same donor

We compared the target specificity of ATHS-C30 on RBD with other mAbs isolated from the same donor (ATHS-C24, ATHS-C34, ATHS-C48, and ATHS-C52) as well as other existing neutralizing mAbs reported previously (THSC20.HVTR04, THSC20-HVTR26, REGN10987, REGN10933) by epitope binning assay as described earlier (27). For this, saturated concentration (100µg/mL) of ATHS-C30 was first incubated with ancestral SARS-CoV-2 RBD bound to Nickel-NTA biosensor, and the binding of the competing mAb at a lower concentration (25µg/mL) to the same RBD was examined. As shown in **Figure 5A**, ATHS-C30 was clearly found to have distinct and non-overlapping epitope specificity on RBD when compared with that of the other four autologous mAbs isolated from A0015 donor, all of which showed additional binding after they were added to RBD bound to a saturated concentration of ATHS-C30. Interestingly, ATHS-C30 demonstrated considerable overlap in target specificity with two existing mAbs, THSC20.HVTR04 (27) and REGN10987 (Imdevimab). However, this observation was not found to be correlated with their neutralization profiles as shown in **Figure 5B**, where unlike ATHS-C30, none of the THSC20.HVTR04 and REGN10987 showed neutralization of XBB1.5 and JN.1 in the pseudovirus neutralization assay. Taken together, our data indicate that ATHS-C30 target epitopes present on the SARS-CoV-2 RBD are evolutionarily conserved in addition to the residues brought in by the SARS-CoV-2 variants through mutations accumulated over time.

**Figure 5.**
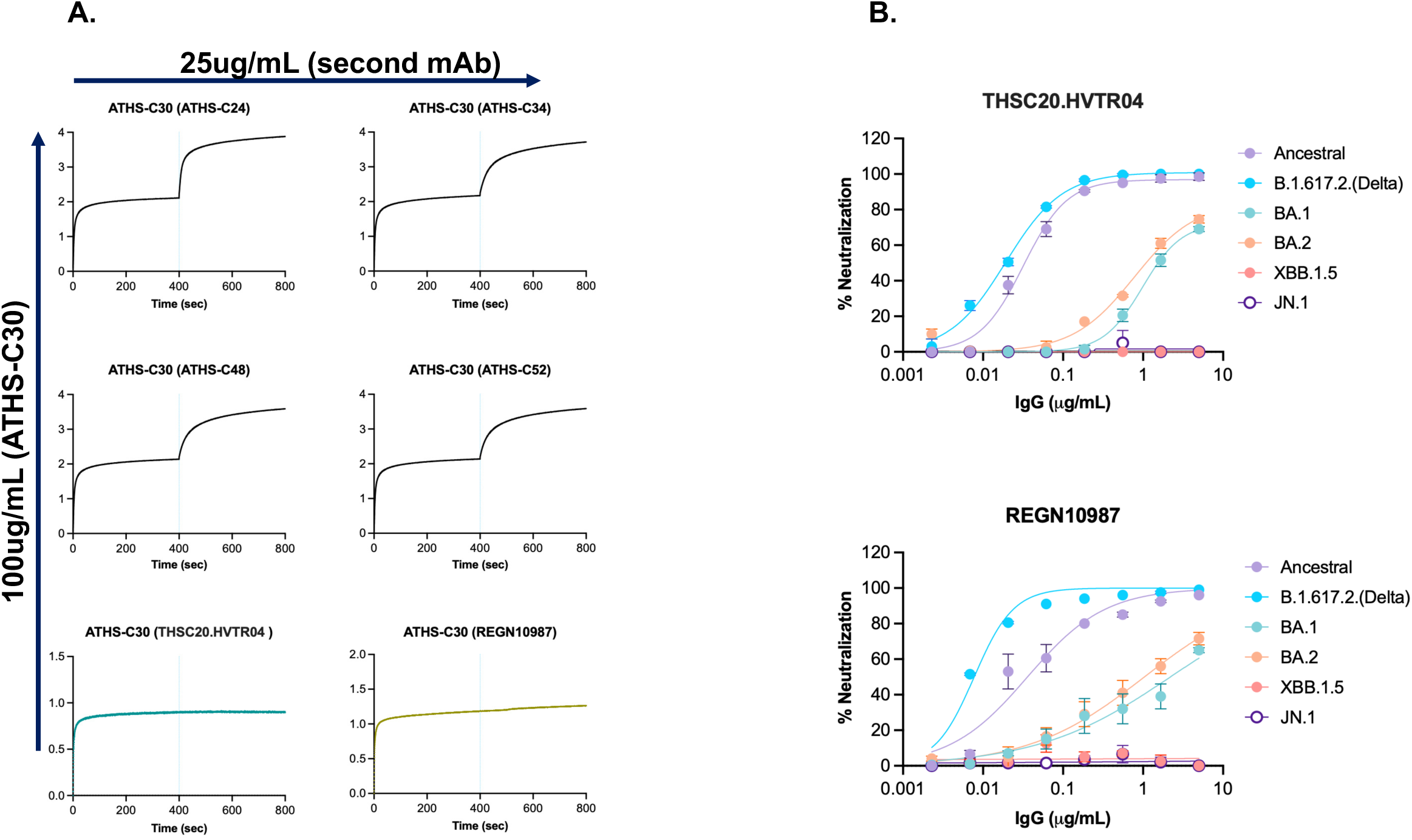
Assessment of overlapping epitope specificity of ATHS-C20 with autologous and heterologous mAbs. **A.** Epitope binning assay using BLI was carried out by first adding an excess amount (100µg/mL) of ATHS-C30 to ancestral biotinylated RBD bound to streptavidin sensor, and subsequently, the second competing mAb (25µg/mL) was added to the reaction. Note that while ATHS-C30 demonstrated distinct epitope specificity on RBD when compared to other mAbs isolated from the same donor, it showed considerable overlapping specificity with that of THSC20.HVTR04 and REGN10987; **B.** Pseudovirus neutralization of XBB.15 and JN.1 by THSC20.HVTR04 and REGN10987 indicated that they differ with ATHS-C30 in their ability to neutralize these two Omicron variants.

### Predicted structural basis of neutralization breadth conferred by ATHS-C30

To elucidate the possible basis for enhanced neutralization breadth conferred by ATHS-C30, a thorough structural analysis was performed using computational methods. The most stable modelled structure of ATHS-C30 was used to generate the complex with JN.1 via docking. The most thermodynamically favorable pose of the complex was used for quantitative analysis to pinpoint the residue level key interactions. We found that ATHSC-30 engages the JN.1-RBD through a network of specific interactions by making hydrogen bonds (HBs) between F377-L:Y92, K378-L:Y92, G404 - L:N94, K440-H:D106, T500-H:54, and H505-H:Y52 (epitope-paratope residue pairs, respectively). Among these interactions, K440-H:D106 is particularly significant, as it is not only involved in making HB but also forms a consistent salt bridge, thereby contributing substantially to the stability of the antibody-antigen complex (**Figure 6**). Furthermore, the epitope conservation analysis demonstrated that the majority of ATHSC-30 contacting residues are evolutionarily conserved across SARS-CoV-2 variants. Some exceptions include residues K440 and H505, which exhibit sequence variability. However, the N440K substitution is unlikely to adversely affect ATHSC-30 binding since both N-to-K substitutions preserve favorable physicochemical characteristics at the interface and lysine remains capable of maintaining stabilizing electrostatic interactions. The overall conservation of the ATHSC-30 epitope indicates a reduced likelihood of immune escape and highlights the potential breadth of ATHSC-30 -neutralization against emerging SARS-CoV-2 variants. Further, to elucidate the possible cause of broad neutralizing nature of ATHSC-C30, the most thermodynamically favorable pose of complex was used for a comparative binding evaluation with reported five different binding classes of neutralizing mAbs against SARS-CoV-2. The binding site comparison with other classes (analyzed using existing structures as shown in **Table S2**) of reported mAbs) indicates that ATHSC-30 belongs to the class 4 category of SARS-CoV-2 neutralizing antibodies (**Figure 7**). The overlay of the ATHSC-30-bound structure with other antibody-RBD complexes showed notable steric clashes between the CDRs of ATHSC-30 and those of THSC20.HVTR04 and REGN10987 indicate that ATHSC-30 shares epitopes that partially overlap with both of these antibodies, consistent with the experimental epitope binning data. The overlap in binding sites likely explains the inability of HVTR04 and REGN10987 to simultaneously bind the RBD in the presence of ATHSC-30 as observed in the epitope binning assay.

**Figure 6.**
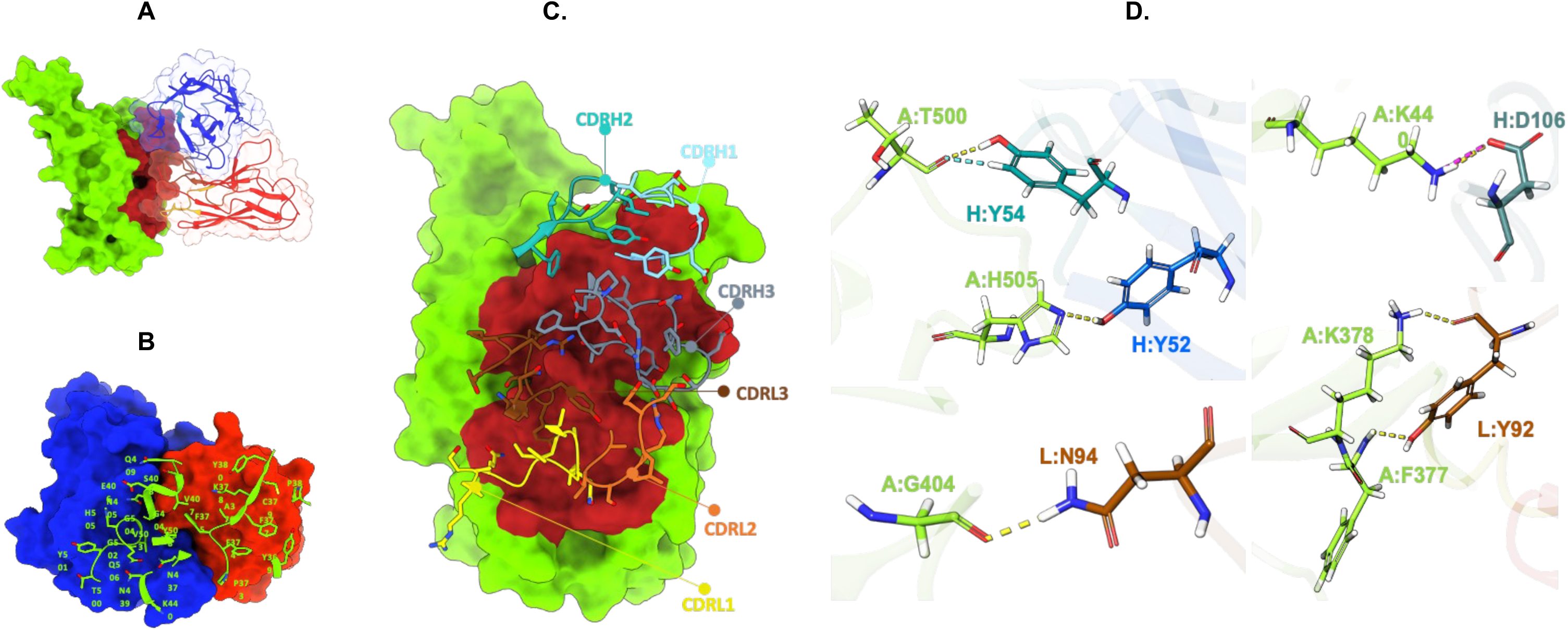
Structural basis of ATHSC-30 recognition of the JN.1 RBD. **A**. Overall binding mode of ATHSC-30 on the JN.1 RBD. The RBD is shown as a green surface, with the ATHSC-30 heavy and light chains represented as blue and red ribbon models, respectively. **B**. Epitope mapping of ATHSC-30 on the JN.1 RBD. Residues comprising the ATHSC-30 epitope are shown as green sticks. The heavy and light chains of ATHSC-30 are represented as blue and red surfaces, respectively. **C.** Paratope analysis of ATHSC-30. The JN.1 RBD is shown as a green surface, while the CDRs of ATHSC-30 are depicted as stick models. **D.** Residue-level interactions between ATHSC-30 and the JN.1 RBD. Representative intermolecular contacts are shown as stick models, with RBD residues colored green and antibody residues colored according to chain identity. Hydrogen bonds are depicted as yellow dashed lines. Key interactions include T500–H:Y54, H505–H:Y52, G404–L:N94, and F377/K378–L:Y92. The interaction between K440 and H:D106 forms both a hydrogen bond and a salt bridge, contributing significantly to stabilization of the ATHSC-30 RBD complex.

**Figure 7.**
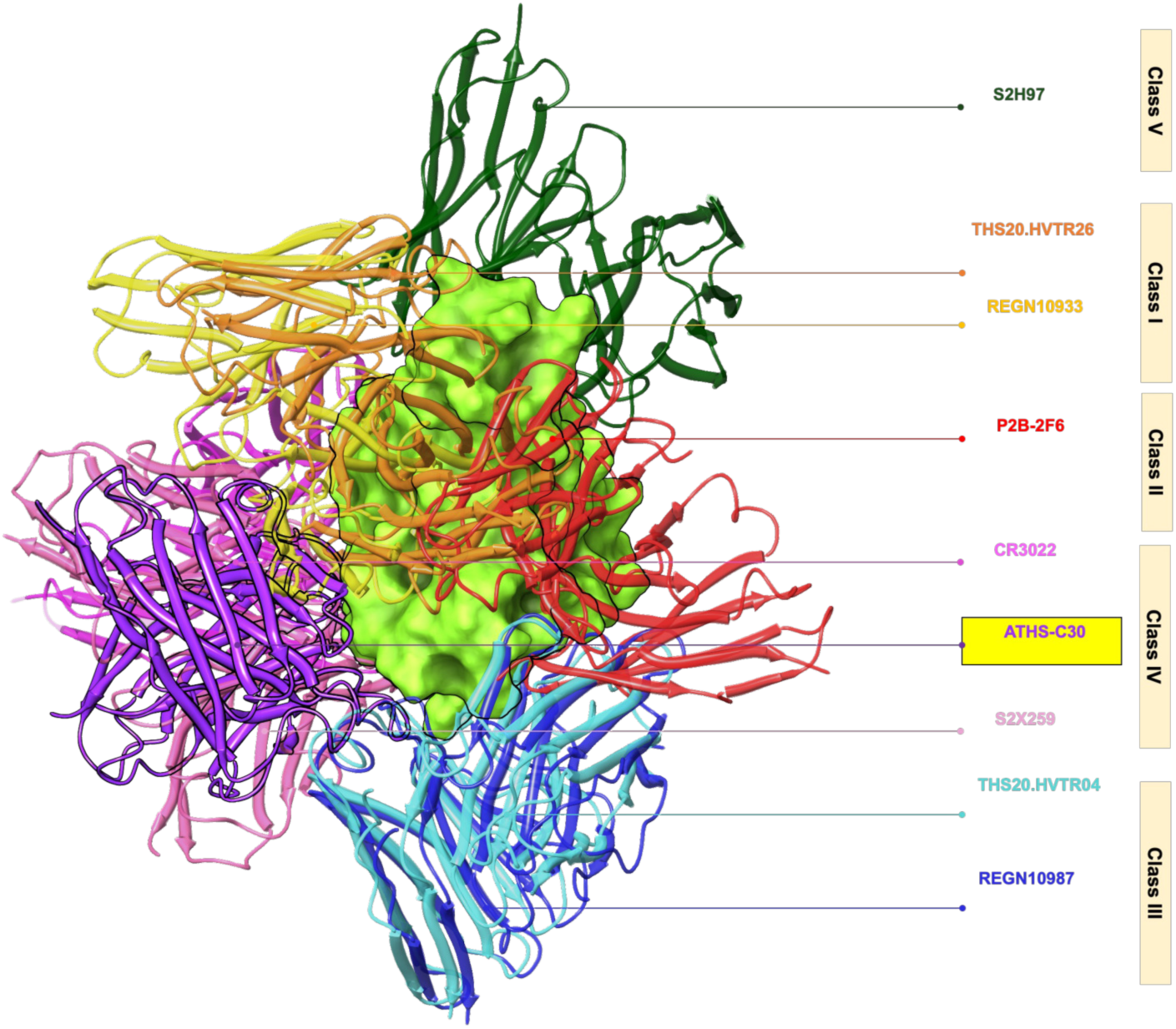
Structural classification of ATHSC-30. Structural superposition of ATHSC-30 with representative antibodies from the five major classes of SARS-CoV-2 RBD targeting antibodies. The RBD is shown as a light green surface, while antibodies are represented as ribbon models: Class I, THS20.HVTR26 (orange; PDB: 7Z0X) and REGN10933 (yellow; PDB: 6XDG); Class II, P2B-2F6 (red; PDB: 8DCC); Class III, THS20.HVTR04 (cyan; PDB: 7Z0Y) and REGN10987 (blue; PDB: 6XDG); Class 4, CR3022 (magenta; PDB: 6ZLR) and S2X259 (pink; PDB: 7M7W); and Class V, S2H97 (dark green; PDB: 7M7W). ATHSC-30 is shown in purple. Structural alignment demonstrates that ATHSC-30 occupies an epitope that overlaps with the binding footprints of the class 4 antibodies CR3022 and S2X259. The observed epitope overlap with established class 4 antibodies indicates that ATHSC-30 recognizes the conserved region of the RBD characteristic of class 4 antibodies. These structural findings classify ATHSC-30 as a Class IV SARS-CoV-2 neutralizing antibody and provide a mechanistic basis for its broad reactivity against antigenically diverse SARS-CoV-2 variants.

## Discussion

Since the beginning of the pandemic in 2019, SARS-CoV-2 evolution has been accompanied by the accumulation of complex mutations, particularly in the ACE-2 receptor-binding domain (RBD) of the viral spike protein, resulting in newly emerging variants capable of evading humoral immune responses either induced by vaccination or mounted through natural infection. Vaccine effectiveness studies have generally shown to provide short-term protection against severe outcomes in infection naïve individuals(12), primarily due to waning of antigen-specific antibodies (28, 29). This formed the basis for introducing a third booster dose, particularly for elderly and vulnerable individuals, to promote the durability of protective antibody levels. SARS-CoV-2 evolution leading to the emergence of Omicron lineages such as BA.5, XBB, JN.1 were accompanied by the accumulation of specific point mutations such as R346T, K417N, L452R, E484A, which showed profound effect on increased transmissibility in one hand and were associated with a significant reduction in neutralization by antibodies induced by vaccination or by prior infection (6, 26, 30). These observations led to several reports that indicated that a booster dose of vaccines or hybrid immunity is a prerequisite for generating robust, qualitatively superior humoral immunity to effectively neutralize Omicron and its lineage variants (13, 14, 31, 32). It is thus believed that the quality of the antibody response developed both through vaccination and in the natural course of infection and reinfection with SARS-CoV-2 determines the extent of protection against severe disease outcomes.

In the present study, we examined the quality, durability and magnitude of a select number of individuals who received three doses of vaccine with a history of infection, thereby categorised as those with hybrid immunity. Most of them, when followed longitudinally, showed durable antibody responses of high magnitude, an observation expected and consistent with previous reports (33, 34). While most of them were found to develop antibodies with specificities to a number of Omicron variants such as BA.1, BA.2.75, BQ.1.1, and XBB.1.5, one individual (A0015) was found to develop antibody responses with extraordinarily expanded specificity to JN.1 and variants of Omicron lineage that emerged subsequently. This observation, indicative of the development of a rare quality of B cell clonotypes in this particular individual, was further examined by isolating mAbs via antigen-specific B cell cloning. All the five functional mAbs isolated from A1005 donor individually and collectively were found to show neutralization breadth including several Omicron variants. Interestingly, one of them (ATHS-C30) was also found to show binding to JN.1 RBD and potently neutralize JN.1 in both pseudovirus and authentic live virus neutralization assays. Furthermore, binding of ATHS-C30 to the JN.1 spike trimers expressed on the cell surface suggested it targets epitopes on the spike protein in its native conformation. To our surprise, we found that ATHS-C30 is able to potently neutralize all the contemporary Omicron variants, including ones that emerged post JN.1 variant such as KP.2, KP.3.1.1, KP.3.2 , KP.3.3, LB.1 and XEC, which suggest it targets evolutionarily conserved residues that serve as targets of antibodies to mediate broad cross-variant neutralization overcoming complex mutational constraints associated with immune escape. One of the striking features observed with ATHS-C30 from the epitope binning experiment carried out using ancestral RBD was that it showed distinct epitope specificity to that of its autologous counterparts isolated from the same donor, as well as a few other existing mAbs. However, when compared with the existing mAbs, THSC20.HVTR04 (27) and REGN10987, we found that ATHS-C30 targets epitopes that overlaps with those targeted by THSC20.HVTR04 and REGN10987 on RBD. Interestingly, despite having overlapping epitope targets on the spike protein, both THSC20.HVTR04 and REGN10987 were found not able to neutralise several contemporary Omicron variants, including JN.1 and its subsequent lineages. These observations indicate that ATHS-C30 possess the following key attributes: (a) it targets residues that are evolutionarily conserved and which form key epitopes for existing broadly neutralising mAbs such as THSC20.HVTR04 and REGN10987, (b) it likely targets non-dominant residues that are brought in by evolving Omicron variants through substitutions or mutations, and (c) it overcomes the mutations such as L455S, K356T, N440K, L452W, Q493E, N501Y, F456L, R346T that are known to confer resistance to existing neutralizing antibodies by JN.1, KP.3.1.1. We believe that specificity to the non-dominant residues provide ATHS-C20 with the ability to confer extraordinary neutralization breadth. The computational aided structural study indicated that Moreover, computational aided structural study predicated that the extraordinary neutralization breadth conferred by ATHSC-30 could be through a mechanism involving destabilization of the spike glycoprotein as few broadly neutralizing class 4 antibodies have been shown to induce conformational rearrangements within the S1 subunit, promoting dissociation of the trimeric spike complex (35, 36). However, this needs to be validated by in-depth experimental structural studies. Because their binding site is highly conserved, class 4 antibodies have been reported (37) to exhibit broad cross-reactivity and effectiveness across multiple sarbecoviruses, including SARS-CoV-1 and SARS-CoV-2 variants. Class 4 antibodies are known to inhibit viral entry by physically preventing the structural shedding of the S1 subunit, that interfere with the fusion of the viral spike with the host cell membrane (38). In conclusion, although hybrid immunity has consistently been associated with the induction of cross-neutralizing antibody responses, the breadth and potency of neutralization against antigenically diverse SARS-CoV-2 variants remain highly heterogeneous across individuals. Consistent with this variability, only donor A0015 in our cohort generated antibodies with exceptional neutralizing breadth, indicating that such responses represent a rare outcome rather than an intrinsic feature of hybrid immunity. While hybrid immunity broadly enhanced the magnitude and durability of the humoral immune response, our findings suggest that the exceptional breadth of ATHS-C30 arose from the recruitment of a rare B-cell clonotype that was primed by the prototype COVID-19 vaccine and subsequently refined through SARS-CoV-2 infection. The resulting class 4 antibody targets an evolutionarily conserved epitope that has remained largely refractory to antigenic drift, thereby retaining potent neutralizing activity against genetically and antigenically diverse SARS-CoV-2 Omicron variants, including contemporary variants that emerged post JN.1 wave. These findings provide evidence that the quality of the B-cell repertoire, rather than hybrid immunity alone, is a critical determinant of the development of exceptionally broadly neutralizing antibodies and highlight the potential of conserved class 4 epitopes as targets for next-generation coronavirus vaccine and immunotherapeutic design.

## Materials and Method

### Ethics statement

Blood samples were obtained from study participants following approval from the Institutional Ethics Committee (IEC) of the All India Institute of Medical Science, New Delhi and also from the IEC of the Translational Health Science & Technology (THSTI), Faridabad, Haryana, India (THS 1.8.1/ (133) dated 15 Nov 2022. Informed consent in English and the local language (Hindi) was obtained from all study participants. The period of recruitment of study participants was between 28/07/2022 and 09/11/2023. Experiments were conducted after approval by the IEC and the THSTI Biosafety (IBSC) committees. Virus isolates from the WHO BioHub were obtained following approvals from the IBSC and the Review Committee on Genetic Manipulation, Department of Biotechnology, Ministry of Science & Technology, Government of India (RCGM).

### Plasmids, cell lines and antibodies

Plasmids used to clone variable heavy and light chains were described earlier (27). pcDNA3.1 plasmid DNA encoding JN.1 full-length spike sequence and 15-mer (GLNDIFEAQKIEWHE) avi-tagged RBD sequence (GISAID accession number EPI_ISL_18644572) were used to produce pseudovirus antigen for neutralization, binding assays and FACS sorting of B cells. pNL4-3.Luc.R-E- backbone was obtained from the NIH AIDS Reagent Reference Program. Human embryonic kidney 293T cells (HEK293T) were used to produce pseudoviruses and recombinant antibodies for screening experiment. Expi-293 and Expi-CHO cells were used to produce and purify small-scale recombinant immunoglobulins (IgGs); HEK293T cells overexpressing ACE-2 (27) (Cat No. 631289, Takara) was used for pseudovirus neutralization assay and Vero-E6 cells were used for live virus neutralization assay. All the cell lines were obtained from the American Type Culture Collection (ATCC).

### RBD-specific B cell sorting

Antigen-specific B cell sorting was carried out as described previously (27). Cryopreserved peripheral blood mononuclear cells (PBMCs) were rapidly thawed at 37°C and resuspended in pre-warmed RPMI 1640 supplemented with 10% fetal bovine serum (FBS). Cell viability and concentration were determined via the trypan blue exclusion method. The avi-tagged SARS-CoV-2RBD antigen was first labelled with biotin (Avidity, BirA500) was subsequently coupled to streptavidin-AF647 (BD Biosciences) by incubating the mixture at room temperature for 45 minutes. Antibody cocktails were prepared in the staining (PBS+0.1%FBS) buffer and incubated for 40 minutes at 4 degrees. After incubation cells were washed twice with FACS buffer to remove unbound antibodies and then filtered through 70-micron filter. For memory B cell, cells were first identified based on light scatter properties (FSC-A vs. SSC-A). Singlet cells were gated based on FSC-H vs. FSC-A gating. Dead cells were excluded using an amine-reactive live-dead stain. PE-Cy7 conjugated CD3^-^, CD8^-^ , CD14^-^, CD16^-^, population excluded as dump channel. Total B cells were gated by BV421conjugated CD19^+^ CD20^+^ and further sub-gated into class-switched PerCP5.5 conjugated IgD^-^, IgM^-^APC H7 conjugated IgG^+^. Further, AF647 conjugated ancestral Wuhan and JN.1 RBD double positive cells were gated. Cells were acquired in FACS symphony analyser. This data analysis was done by using Flow jo v10 software.

### Amplification and cloning of variable heavy and light IgG chains

cDNA was prepared using Superscript III Reverse Transcription kit from sorted B cells as described before (27). cDNA master mix containing dNTPs, random hexamers, IgG gene-specific primers and RT enzyme were added to generate cDNA. Heavy and light-chain variable regions of IgG were amplified in independent nested PCR reactions using gene specific primers. First round PCR amplification was performed using HotStar Taq DNA Polymerases (Qiagen) and second round nested PCR was performed using Phusion HF DNA polymerase (Thermo Fisher Inc.). Specific restriction enzyme cutting sites (heavy chain, 5′-AgeI/3′-SalI; kappa chain, 5′-AgeI/3′-BsiWI; and lambda chain, 5′-AgeI/3′-XhoI) were introduced in the second round PCR primers to clone into the respective expression vectors. Amplified PCR products were verified on the agarose gel and wells with double positives (with amplification of both Heavy and Light chain variable regions from the same well) were identified and selected for subsequent cloning experiments. PCR products were digested with specific restriction enzymes, purified and cloned in-frame into expression vectors encoding the human IgG1, Ig kappa or Ig lambda constant domains using the Quick Ligase cloning system (New England BioLabs) according to the manufacturer instructions. Ligation reactions were transformed into NEB 5-alpha competent *E*. *coli* cells, plated on LB agar plates containing ampicillin and incubated overnight at 37 °C. Colonies with the desired inserts were screened by colony PCR and inoculated to prepare plasmid DNA. Plasmid clones with the desired inserts were further confirmed by restriction digestion with respective restriction enzymes (New England Biolabs, Inc.). Confirmed heavy and light chain plasmid DNAs were co-transfected in HEK293T cells using Polyethylene imine (PEI)in 24-well tissue culture plates for preparing recombinant antibody supernatant for initial screening for their expression, antigen specificity and neutralization ability.

### Expression of recombinant mAbs

Recombinant human monoclonal antibodies (mAbs) were produced using the HEK293T cell expression system. Cells were maintained in Dulbecco’s Modified Eagle’s Medium (DMEM) supplemented with 10% heat-inactivated fetal bovine serum (FBS), 2 mM L-glutamine, and 1X antibiotic-antimycotic at 37°C in a 5% CO_2_ atmosphere. For transient transfection, cells were seeded in 24-well plates and grown to 80% confluency. Heavy (IgH) and light (IgL) chain expression vectors (750 ng each) were complexed with polyethyleneimine (PEI, 4.5 µg) in 100 µL of Opti-MEM (Invitrogen Inc.). After a 20-minute incubation at room temperature, the DNA-PEI complexes were added dropwise to the cultures and transfected cells were incubated for 72 hours. The antibody-containing supernatants were harvested by centrifugation at 800 x g for 10 minutes and stored at 4°C for downstream functional characterization.

### IgG purification

Clarified cell culture supernatants collected post-transfection into suspension cell line Expi293/ExpiCHO cells using PEI transfection reagent with up to five days incubation at 37^0^ C in a humidified CO_2_ shaker incubator) were filtered through 0.22 µm membranes and incubated overnight at 4°C with Protein A resin (G-Biosciences Inc.; 500 µl resin per 100 mL supernatant) under constant agitation. The resin was packed into a chromatography column and washed extensively with 1 M Tris-HCl buffer to remove non-specific contaminants. Bound IgG was eluted using 100 mM glycine-HCl (pH 2.5) directly into tubes pre-equilibrated with 1M Tris-HCl (pH 8.0) to ensure immediate neutralization and maintain protein stability. The eluted fractions were then transferred to 30 kDa molecular weight cut-off (MWCO) dialysis membranes for overnight buffer exchange into PBS at 4°C. Post-dialysis, the antibodies were concentrated using 10 kDa Amicon centrifugal filter units. Final IgG concentrations were determined by measuring absorbance with a Nanodrop spectrophotometer.

### Quantification of Plasma Anti-SARS-CoV-2 RBD-Specific IgG by ELISA

Plasma anti-SARS-CoV-2 receptor-binding domain (RBD)-specific IgG levels were determined using the in-house RBD-based IgG ELISA as described previously (39). The assay used recombinant RBD from the ancestral strain of SARS-CoV-2 as the target antigen. Based on the World Health Organization (WHO) manual for the establishment of national and other secondary standards for antibodies against infectious agents, the kit was calibrated against the First WHO International Standard for anti-SARS-CoV-2 immunoglobulin, NIBSC code 20/136. Anti-RBD IgG concentrations were expressed as Binding Antibody Units per millilitre (BAU/mL). An anti-RBD antibody titre greater than 24 BAU/mL was considered positive.

### RBD binding assay by ELISA

The binding of purified and unpurified antibodies to SARS-CoV-2 spike’s RBD and Omicron JN.1 spike’s RBD was determined using an ELISA assay. RBD was coated onto the ELISA plate at a concentration of 2µg/mL in PBS solution, using a 50 µL volume for overnight incubation at 4°C. Following incubation, the plates were washed three times with 200 µL of PBST (PBS containing 0.05% Tween-20) and blocked with 100 µL of 5% (w/v) skimmed milk for 1 hour at 37 degrees to prevent nonspecific binding. After blocking, the plates were washed three times with PBST and 50 µL of each sample supernatant was added to the respective wells. The plates were incubated for 1 hour at 37°C, followed by three washes with PBST to remove unbound antibodies. Subsequently, 50 µL of horseradish peroxidase (HRP)- conjugated goat anti-human IgG (diluted 1:4000 in 2.5 % skimmed milk) was added to each well and incubated for 1 hour at 37°C. Color development was achieved by adding TMB (3,3’,5,5’-tetramethylbenzidine) substrate and the enzymatic reaction was stopped by adding 1 N HCl. The optical density (OD) was measured at 450 nm using a multimode plate reader .

### Pseudovirus neutralization assay

For the pseudovirus production, HEK-293T cells were seeded in 6-well plates at density of 4 X 10^6^ cells per plate with 2 mL of DMEM supplemented with 10 % FBS. Upon reaching approximately 80% confluency, the cells were transfected with SARS-CoV-2 or Omicron JN.1 spike expression plasmid DNA, together with pNL4-3.Luc.R-E- used as a backbone (40) using polyethylenimine (PEI). The plasmid DNA and PEI were mixed Opti-MEM medium and incubated for 20 minutes at room temperature to allow complex formation. Subsequently, 200 µL of DNA-PEI complex was added dropwise to each well and incubated for 72 hours at 37°C. Following incubation, the supernatant containing pseudovirus particles were collected and centrifuged at 700rpm for 10 minutes to remove cellular debris. For neutralization assay, 50 µL of pseudovirus was mixed with 50 µL of antibody-containing supernatant or purified IgG and incubated at 37°C for 1 hour to allow antibody-virus complex formation. After incubation, 100 µL of ACE-2 expressing HEK293T target cells of density 10,000 cells/well supplemented with DEAE-dextran (Diethylaminoethyl dextran) were added to each well and the plates were incubated for 72 hours at 37°C. Post incubation, 70ul of the culture medium was removed from each well and 40 µL of Britelite substrate was added and incubated for 2 minutes at room temperature. Luminescence was measured using a luminometer and the relative lights units (RLU) were recorded to assess the pseudovirus infection and neutralization efficiency. Neutralization curves were generated, and IC₅₀ values were calculated in GraphPad Prism using a non-linear regression model (log[inhibitor] versus normalized response, variable slope 4 parameters).

### Live virus neutralization assay

Vero-E6 cells were cultured in Minimal Essential Medium (MEM) supplemented with 10% heat inactivated fetal bovine serum (FBS), 1% penicillin-streptomycin and 1% non-essential amino acid. Cells were seeded at a density of 25,000 cells per well in 96-well plates and incubated overnight at 37°C with 5% CO₂. Antibody reagents were prepared as two-fold dilutions in dilution medium (MEM containing 1% FBS), ranging from 5 to 0.00244μg/mL (75μL reagent + 75μL diluent per step). An equal volume (75 μL) of pre-titrated SARS-CoV-2, BA.5, XBB.1.16, JN.1, KP.2, KP3.1.1, KP.2, KP.3.1.1, KP.3.2, KP3.3, LB.1 and XEC virus suspension—calibrated to yield approximately 60–200 foci in virus-only controls—was added to each dilution and incubated for 1 h at 37°C with 5% CO₂. A predefined SARS-CoV-2 neutralizing antibody-positive serum was included as an assay control. Following the neutralization step, 100μL of the virus–reagent mixture was transferred onto Vero E6 monolayers and incubated for 1 h at 37°C with 5% CO₂ to allow virus adsorption. After adsorption, an overlay medium was added, and plates were incubated for 24 h. For Omicron sub-lineages, incubation was extended to 28 h. Cells were then fixed and permeabilized with 100 μL of IMF permeabilization buffer at room temperature for 20 min. Viral foci were detected by immunostaining with anti-nucleocapsid primary antibody for 1h, followed by staining with Alexa Fluor 488–conjugated donkey anti-mouse IgG secondary antibody (1:500 dilution). After three washes with Milli-Q water, foci were quantified using a CTL Immuno-Spot Reader. Neutralization curves were generated, and IC₅₀ values were calculated in GraphPad Prism using a non-linear regression model (log[inhibitor] versus normalized response, variable slope).

### BLI kinetics

Binding affinities of selected monoclonal antibody (mAb) to SARS-CoV-2 and Omicron JN.1 spike receptor binding domains (RBDs) were measured using biolayer interferometry (BLI) on an Gator Prime platform at 30°C with agitation at 1,000 rpm. Ni-NTA biosensors were equilibrated in 1X PBS solution for 15 min prior to use. Ancestral SARS-CoV-2 or Omicron JN.1 RBD or spike was captured onto the sensors to a loading response of ∼1.0 nm. Loaded sensors were briefly equilibrated in PBS for 60 s before being transferred into wells containing mAb at defined concentrations to monitor association for 400 seconds. A five-point, three-fold serial dilution series of each mAb (10, 3.3, 1.1, 0.36, and 0.12 µg/mL) was prepared in PBS with 0.02% Tween 20. Following the association, sensors were immersed in PBS for 400 s to record dissociation. Reference subtraction and alignment to the baseline were performed using Gator Prime data Analysis Software. Binding curves were globally fitted using a 1:1 interaction model, and kinetic parameters (Kon, Koff) as well as equilibrium dissociation constants (K_D_) were calculated from the global fit.

### Epitope binning assay

Epitope binning analyses were conducted using Streptavidin (SA) and Protein A biosensors. . All incubations were performed in 1× PBS + 0.02% Tween 20. Biotinylated RBD antigens (10 µg/mL, 200 µL) were immobilized on SA biosensors, or mAb (10 µg/mL) was loaded onto Protein A biosensors, to achieve a loading response of approximately 0.9–1.3 nm, followed by a 60-s wash in PBS. Sensors were then immersed in saturating concentrations of the primary mAb (100 µg/mL) for 500 s to allow full binding. Subsequently, sensors were transferred to wells containing the competing secondary mAb at 25 µg/mL for 400s to assess additional binding in the presence of the saturating antibody.

### Cell surface spike binding assay

The binding of monoclonal antibodies to SARS-CoV-2, Omicron HV.1, XBB.1.5 and JN.1 spike variants expressed on the surface of HEK293T cells was evaluated by flow cytometry. HEK293T cells were transiently transfected with the spike-expressing plasmid DNA and PNL4.3 backbone and incubated for 36-48 hour at 37°C in 6 well plate. Following the expression, cells were washed with PBS and trypsinized and neutralized by adding complete media (10% FBS+DMEM). Cells were pelleted down at 1500 RPM for 5 minutes at 4 degrees. Resuspended in FACS buffer and 100,000 cells per well were dispensed into 96-well U-bottom plates. Threefold dilution of mAb were prepared starting at 5 μg/mL and extending to 0.041 5 μg/mL in a total volume of 50μL per well. These dilutions were added to spike-expressing and untransfected control cells, which were then incubated on ice for 1 hour. After incubation, cells were washed twice with FACS buffer (1 X PBS, 2% FBS, 1mM EDTA) and stained with a 1:200 dilution of R-Phycoerythrin (PE)–conjugated Affini-Pure F(ab’)₂ Fragment Goat Anti-Human IgG, F(ab’)₂ fragment-specific antibody (Jackson Immuno Research) for 30 min. Cells were further stained with LIVE/DEAD Fixable Aqua viability dye (Thermo-Fisher) for 15 min, washed twice with FACS buffer, and analysed on a BD FACS Canto flow cytometer. The percentage of PE-positive cells was quantified to assess mAb binding. A mock infection was used as a negative control. Data analysis was performed using FlowJo 10 v software.

### Sanger sequencing

Plasmid DNA from selected antibody-expressing clones was isolated using a mini-prep protocol according to the manufacturer’s protocol. The concentration of plasmid was measured using a spectrophotometer. Plasmid DNA was sequenced by Sanger sequencing method. Sequencing reactions were prepared using either specific forward or reverse primers. Sequencing was performed by an external commercial facility platform. Raw chromatogram files were inspected manually using sequence analysis software (BioEdit) to ensure high-quality base calling and clean sequences. The verified sequence was then loaded into IMGT IgG BLAST tool for further analysis.

### Computational studies

The protein structures of the SARS JN.1 (PDB ID:9LOY) (36) and antibodies were curated from the RCSB in the PDB format (http://www.rcsb.org/pdb). For JN.1 only receptor binding domain (RBD) was used, rest of the spike protein was deleted. These structures, along with their corresponding resolution values, are summarized in **Table S2**. All structures were prepared using Maestro’s Protein Preparation Wizard module (Schrödinger Release 2025-4). The structures were curated by removing spike proteins, water molecules, and metal ions. Hydrogens and bond orders were added, and missing side chains and loops were filled in using Prime. Hydrogen bond (HB) optimization and restrained minimization were performed for the systems using force field OPLS4 (41).

### Molecular modelling

A15C30 was modelled from the sequence of heavy and light chain identified from Sanger sequencing. Two approaches were used to generate the 3D model of A15C30 i.e Homology modelling (using Biologics module within the Schrödinger Release 2025-4 software) and *de-novo* modelling (using AF3) (**Figure S2**). Both systems were subjected to 500ns MD simulations to allow the modelled structure to attain its stable conformation (**Figure S3**). Furthermore, the PCA analysis was performed, which led to diverse conformations for CDR regions obtained from homology and from a *de novo* model (**Figure S4 and S5**). All the structures were similar except at the CDR region where conformational changes were observed.

### Protein-antibody complex formation via docking followed by MD simulation

The modeled antibody structure (via homology and using Alpha fold 3) was then docked to the JN.1 RBD using PIPER module of Schrodinger. The non-CDR regions of the antibody were masked during the process. The CDR region was defined according to the Chothia definition (42). The best docked complex was used for MD simulation (100ns) to achieve the most native like stable structure. The stable frame from the simulation (lowest minima) was used for interaction analysis and for comparison with co-crystals.

## Acknowledgments.

We thank the study volunteers and all the collaborators for their support. This research was conducted using biological materials: JN.1/Omicron, KP.2.16, KP.3.1.1, KP.3.2., KP.3.3, LB.1, XEC isolates, obtained through the WHO BioHub System, from the WHO BioHub Facility: Spiez Laboratory, Switzerland. We thank Deepak Rathore for assistance with FACS sorting, the Bioassay laboratory for facilitating the propagation of SARS-CoV-2 isolates, and the THSTI monoclonal antibody biofoundry facility for facilitating the analytical testing of the purified monoclonal antibody. DR was supported by a Senior Research Fellowship from the Department of Biotechnology (DBT/2021-22/THSTI/1516). SD, SA and NK are supported by the Translational Research Program, supported by the Department of Biotechnology.

## Supporting files

**Table S1.** Booster vaccination, gender distribution and sample collection time of individuals with hybrid immunity studied.

**Table S2. References of structures and their resolutions.**

**Figure S1. Isolation of RBD-specific monoclonal antibodies. (A).** Antigen-specific single B cell sorting was done using the 1 x 10^6^ million PBMCs obtained from A0015 donor. Ancestral and JN.1 RBD^+^/IgG^+^/CD19^+^/CD20^+^/IgD^-^/IgM^-^ cells were single sorted in 96-well trays and lysed cells were used as source of RNA for amplification of variable heavy and light chain IgG genes by RT-PCR as described before (27). **B.** Screening for cell-supernatants expressing SARS-CoV-2 RBD-specific recombinant mAb clones. Neat supernatants harvested from 293T cells co-transfected with plasmids expressing variable heavy and light chain IgGs were assessed for their ability to bind to both ancestral and JN.1 by ELISA to down-select antigen-specific functional mAb clones that were subsequently examined for their neutralization potential as shown in Table 1.

**Figure S2.** Workflow for structural modelling and conformational analysis of ATHSC-30.

**Figure S3. Structural stability assessment of de novo and homology models during molecular dynamics simulations.** RMSD profiles of the backbone atoms for the de novo model (orange) and homology model (blue) over a 500 ns molecular dynamics simulation. RMSD values were calculated relative to the respective starting structures and are plotted as a function of simulation time. Both models exhibit an initial equilibration phase followed by convergence to stable conformational ensembles. The de novo model maintains lower RMSD values throughout the simulation, fluctuating predominantly between 1.0 and 1.8 Å, indicating greater structural stability and reduced conformational drift. In contrast, the homology model shows larger deviations, with RMSD values ranging from 2.0 to 2.8 Å. At around 325 ns of the simulation, there was enhanced structural flexibility and sampling of a broader conformational space.

**Figure S4. Homology modelling and principal component analysis. A.** Free-energy landscape projected onto the first two principal components (PC1 and PC2). The contour map depicts the conformational space explored during the simulation, with color intensity corresponding to normalized probability density. Two major low-energy basins, designated Min1 and Min2, represent the most populated and thermodynamically favorable conformational states. **B.** Histograms of projections along PC1 (left) and PC2 (right) reveal the dominant modes of motion captured during the simulation. **C.** Representative conformational states identified from PCA of the 500ns molecular dynamics trajectory. The structures corresponding to the centers of the two dominant free-energy minima are shown as ribbon representations and superimposed onto each other. Two conformations at Min1 and Min2 is colored blue (Heavy chain light blue light chain dark blue) and green (Heavy chain light green light chain dark green) respectively. Structural differences between the two states are primarily observed in the flexible loop regions, indicating distinct conformational substates sampled during the simulation.

**Figure S5. De novo modelling and principal component analysis. A.** Free-energy landscape projected onto the first two principal components (PC1 and PC2). The contour map depicts the conformational space explored during the simulation, with color intensity corresponding to normalized probability density. Three major low-energy basins, designated Min1, Min2 and Min3, represent the most populated and thermodynamically favorable conformational states. **B.** Histograms of projections along PC1 (left) and PC2 (right) reveal the dominant modes of motion captured during the simulation. **C.** Representative conformational states identified from PCA of the 500ns molecular dynamics trajectory. The structures corresponding to the centers of the two dominant free-energy minima are shown as ribbon representations and superimposed onto each other. Three conformations at Min1, Min2 and Min3 is colored green (Heavy chain green light chain olive green), Pink (Heavy chain light pink light chain pink), Purple (Heavy chain magenta light chain purple) respectively. Structural differences between the three states are primarily observed in the flexible loop regions, indicating distinct conformational substates sampled during the simulation. Differences in three conformations were less when compared with homology models.

